# Matrix Metalloproteinase Activity During Methamphetamine Cued Relapse

**DOI:** 10.1101/2022.10.28.514236

**Authors:** Stacia I. Lewandowski, Ritchy Hodebourg, Samuel K. Wood, Jordan S. Carter, Katherine Nelson, Peter W. Kalivas, Carmela M. Reichel

## Abstract

Relapse to drug seeking involves transient synaptic remodeling that occurs in response to drug associated cues. This remodeling includes activation of matrix metalloproteinases (MMPs) to initiate catalytic signaling in the extracellular matrix (ECM) in the nucleus accumbens core (NAcore). We hypothesized that MMP activity would be increased in the NAcore during cue-induced methamphetamine (meth) seeking in a rat model of meth use and relapse. Male and female rats had indwelling jugular catheters and bilateral intracranial cannula targeting the NAcore surgically implanted. Following recovery, rats underwent meth or saline self-administration (6hr/day for 15 days) in which active lever responding was paired with a light+tone stimulus complex, followed by home cage abstinence. Testing occurred after 7 or 30 days of abstinence. On test day, rats were microinjected with a FITC-quenched gelatin substrate that fluoresces following cleavage by MMPs, allowing for the quantification of gelatinase activity by MMP-2 and −9 during cued relapse testing. MMP-2,9 activity was significantly increased in the NAcore by meth cues presentation after 7 and 30 days of abstinence, indicating that remodeling by MMPs occurs during presentation of meth associated cues. Surprisingly, while cue-induced seeking increased between days 7 and 30, suggesting behavioral incubation, MMP-2,9 activity did not increase. These findings indicate that while MMP activation is elicited during meth cue-induced seeking, MMP activation did not parallel the behavioral incubation that occurs during extended drug abstinence.

## 1. Introduction

Methamphetamine (meth) is a widely abused psychostimulant and meth use disorder (MUD) is a chronic relapsing brain disease characterized by compulsive drug seeking and drug use with no FDA approved pharmacotherapies. Drug cravings during abstinence can persist for months and years, which results in a relapse rate of 61% in people suffering MUD during their first year of abstinence^1^. Therefore, it is imperative to explore novel cellular and molecular mechanisms that are associated with and may underpin meth craving to discover therapeutics to treat MUD and prevent relapse.

In rats, incubation of meth craving can occur during forced home cage abstinence from meth self-administration causing cue-induced meth seeking to progressively increase over the course of withdrawal^2,3^. Relapse to meth use triggered by exposure to drug-associated cues recruits multiple brain regions^4^, particularly involving the nucleus accumbens core (NAcore)^5–7^ and dorsal striatum (DS)^8–10^.

Importantly, chronic meth use leads to cognitive deficits that may exacerbate drug relapse^11–13^. It is hypothesized that these cognitive deficits may be a consequence of the neurotoxicity produced by chronic meth use^14,15^. In addition, meth causes blood brain barrier (BBB) dysfunction via reduced expression of structural tight junction proteins^16^ and gap junction proteins^17^. Increased BBB permeability resulting from meth use can lead to neuroinflammation and activation of matrix metalloproteinases (MMPs), including MMP-2 and MMP-9^16^. These proteinases are extracellular matrix (ECM) signaling proteins and are implicated in degradation of the BBB due to meth-induced neuroinflammation^18,19^. MMP-2 and MMP-9 are also integral signaling proteins in the ECM and are recruited during events of synaptic plasticity, such as learning and memory^20,21^. The primary function of the ECM is to maintain structural support of synapses, and is made up of proteins, polysaccharides, perineuronal nets and cell-adhesion molecules. The pre- and post-synaptic neuronal terminals, the ECM, and astroglia together make up what is referred to as the tetrapartite synapse^22,23^. MMP-2 and MMP-9 constitute the gelatinase subfamily and regulate synaptic plasticity via degradation of ECM glycoproteins, allowing for receptor trafficking and synaptic remodeling in the tetrapartite synapse following exposure to drugs of abuse^21,24^. Important for the context of this manuscript, drug induced synaptic remodeling of the ECM by MMPs occurs in the NAcore during drug seeking after cocaine, nicotine, and heroin cue-induced reinstatement^25,26^. MMP activity has not been investigated in an abstinence-relapse model of self-administration during cued meth seeking. Here, we evaluate the hypotheses that MMP’s are activated during relapse to meth-associated cues and that MMP activity will increase concurrent with the incubation of meth seeking.

## 2. Materials and Methods

### 2.1 Animals, Housing, Surgery

Aged-matched male (250-275g) and female (225-250g) Long-Evans rats (Envigo, Indianapolis, IN, USA) were double housed in a climate-controlled vivarium on a reversed 12:12 light/dark cycle. All experiments occurred during the dark phase. Food was restricted to 20g chow/day during experimentation and water was available *ad libitum*. Animal procedures were compliant with the National Institutes of Health Guide for the Care and Use of Laboratory Animals. Rats were anesthetized with isoflurane and anesthesia was maintained at 2% isoflurane. Ketorolac (2.0mg/kg, i.p.) and cefazolin (200mg/kg, s.c.) were given perioperatively for analgesia and prevention of post-surgical infection, respectively. Indwelling jugular catheters were surgically implanted as described previously ^27^. All rats were implanted with intracranial cannulas for *in vivo* zymography. Singular, bilateral guide cannulas (P1 Technologies, Roanoke, VA) were directed at the NAcore (anterior-posterior [AP] +1.6mm, medial-lateral [ML] +/-1.5mm, dorsal-ventral [DV] −5mm, with an injection cannula extending 2 mm beyond the guide for a final DV depth of 7.5mm) and affixed with dental acrylic.

### 2.2 Drugs and Reagents

Methamphetamine HCL (National Institute on Drug Abuse, Bethesda, MD) was dissolved in sterile saline. FITC-Gelatinase (Thermo Fisher Scientific, Waltham, MA; #D12060) was used for *in vivo* zymography at a concentration of 1mg/mL in phosphate-buffered saline (PBS) and was microinjected at 1.5 μL/hemisphere.

### 2.3 Self-Administration

Rats underwent daily 6-hour methamphetamine or saline self-administration sessions in two-lever operant chambers (30 × 20 × 20 cm, Med Associates, Fairfax, VT) as previously described ^27^. Active lever presses were paired with cues (light and tone) and delivered a 2-s infusion of meth (0.4mg/mL) and followed by a 20-second timeout period. Rats self-administered methamphetamine for 15 days, maintaining criteria of 10 infusions a day. Following the last day of self-administration, rats underwent home cage abstinence for 7 or 30 days to model early and late abstinence^28^. Tissue was collected 24-hours after the last day of abstinence either from their home cage (withdrawn group) or a cued-relapse test (30 minutes).

### 2.4 Cue Relapse Testing

Cue relapse testing occurred by returning rats to the operant chambers after a period of home cage abstinence (either 7 or 30 days). No extinction training occurred. FITC-Gelatinase (Thermo Fisher Scientific, Waltham, MA; #D12060; 1mg/ML PBS, pH 7.2-7.4, 1.5μL per hemisphere) was microinjected into the NAcore, rats were returned to the home cage for 15 mins and then placed into the chamber for a 30 min test. Withdrawn rats were not placed in a chamber but were sacrificed at this time. Behaving rats were sacrificed after the test session. A single non-contingent cue presentation signaled the start of the 30-minute test. Rats were not tethered and did not receive any meth or saline. Active lever responding resulted in presentation of the light-tone stimulus that was previously paired with meth during self-administration. Inactive lever responding was without consequence.

### 2.5 In Vivo Zymography, Confocal Microscopy and Quantification

Activity of MMP-2,9 was assessed via in vivo zymography. Rats were transcardially perfused with 0.1M Phosphate Buffer (PB) and followed by 4% formalin. 100μM thick brain slices from the NAcore were obtained from a Vibratome. Only slices containing the anterior commissure, a landmark for the NAcore, and NAcore tissue containing the cannula tract was imaged and analyzed (Image J Version 2, National Institute of Mental Health, Bethesda, MD). The tissue slice with the highest integrated density value was quantified from each animal^25,26^. Z-stacks were obtained via a STELLARIS Leica SP5 laser scanning confocal microscope (Leica, Wetzlar, Germany). MMP integrated density was quantified using ImageJ software.

### 2.6 Statistical Analysis

Statistical analyses were conducted using GraphPad Prism version 9 (GraphPad Software, San Diego, CA). Both sexes were included in these studies as it has been demonstrated that there are sex differences regarding acquisition meth self-administration^13,29^. However, we did not find evidence of sex differences in behavioral relapse testing or MMP activity and thus data are collapsed across both sexes. The means and standard errors are desegregated by sex in Table 1.

**Table 1.** Mean and standard errors desegregated by sex.

|  | Females |  | Males |  |
| --- | --- | --- | --- | --- |
|  | Saline | Meth | Saline | Meth |
| **Day 7** |  |  |  |  |
| Active Lever | 1.333 ± 0.67 | 26.38 ± 6.18 | 3.5 ± 1.19 | 27.00 ± 9.85 |
| Inactive Lever | 1.333 ± 0.88 | 10.5 ± 1.62 | 3.25 ± 1.49 | 4.00 ± 3.51 |
| MMP-2,9 ID | 4265343324<br>± 2984891451 | 15019642860<br>± 3223805646 | 7935662010<br>± 4045981630 | 7896444764<br>± 3052593894 |
| **Day 30** |  |  |  |  |
| Active Lever | 10.00 ± 3.000 | 52.14 ± 11.39 | 5.500 ± 5.500 | 35.00 ± 12.20 |
| Inactive Lever | 5.500 ± 1.500 | 8.714 ± 3.068 | 3.500 ± 3.500 | 8.750 ± 8.097 |
| MMP-2,9 ID | 571957932 ±<br>87232202 | 7613373395 ±<br>927666319 | 2042673410 ±<br>1394526319 | 2494160725 ±<br>1922862467 |

Active and inactive lever responding from acquisition of self-administration was analyzed via a mixed-effects three-way analysis of variance (ANOVA) with drug (meth or saline), sex (male or female) as the between subjects independent variables and time (days 1-15) as the repeated measure. Intake (mg/kg) for meth and saline was analyzed via a two-way ANOVA with sex (males or female) and day (days 1-15) as the repeated measure. Active and inactive lever responding during cued relapse was analyzed with a mixed effects two-way ANOVA with lever (active and inactive) as the within subjects’ variable and drug group as the between subjects variable with Holm-Sidak’s post doc tests. The saline cue groups from 7 and 30 abstinence days were pooled due to no differences between them resulting in 3 drug groups Saline Cue, Meth Cue (day 7), and Meth Cue (day 30).

Integrated density for MMP activity included the saline and meth withdrawn groups and was analyzed via a one-way ANOVA and Holm-Sidak’s post-hoc tests. Five statistical outliers were removed after identification with ROUT (Q=1%) test. One outlier was from the Saline withdrawn and Meth withdrawn groups each, and 3 were from the Saline cue group. Three animals had chewed cannulas becoming unable to receive a microinjection were placed into the withdrawn group (n=3). In all cases, *p* values <0.05 were considered significant.

## 3.0 Results

### 3.1 Rats Acquire Stable Methamphetamine Self-Administration

Rats were trained to self-administer saline or meth for 15 days and were withdrawn (home cage abstinence) for 7 or 30 days prior to cued relapse testing (see Fig 1A for timeline). Meth rats increased active lever pressing (Fig 1B) over time, whereas saline rats did not [day x drug interaction, *F* (14, 784) = 2.99, *p* < 0.0002; and main effect of drug, *F* (1, 784) = 3.148.1, *p* < 0.0001]. Meth males (n=18) had higher lever pressing than females (n=19) [day x sex interaction, *F* (1, 784) = 10.81, *p* < 0.0011; and main effect of sex *F*(1,799) = 150.3, *p* < 0.0001]. The 3-way interaction was not significant [day x sex x time interaction, *F* (14, 784) = 0.724, *p* = 0.75]. There were no differences between drug group, sex, or day on the inactive lever nor were there any significant interactions (Fig 1C).

**Figure 1.**
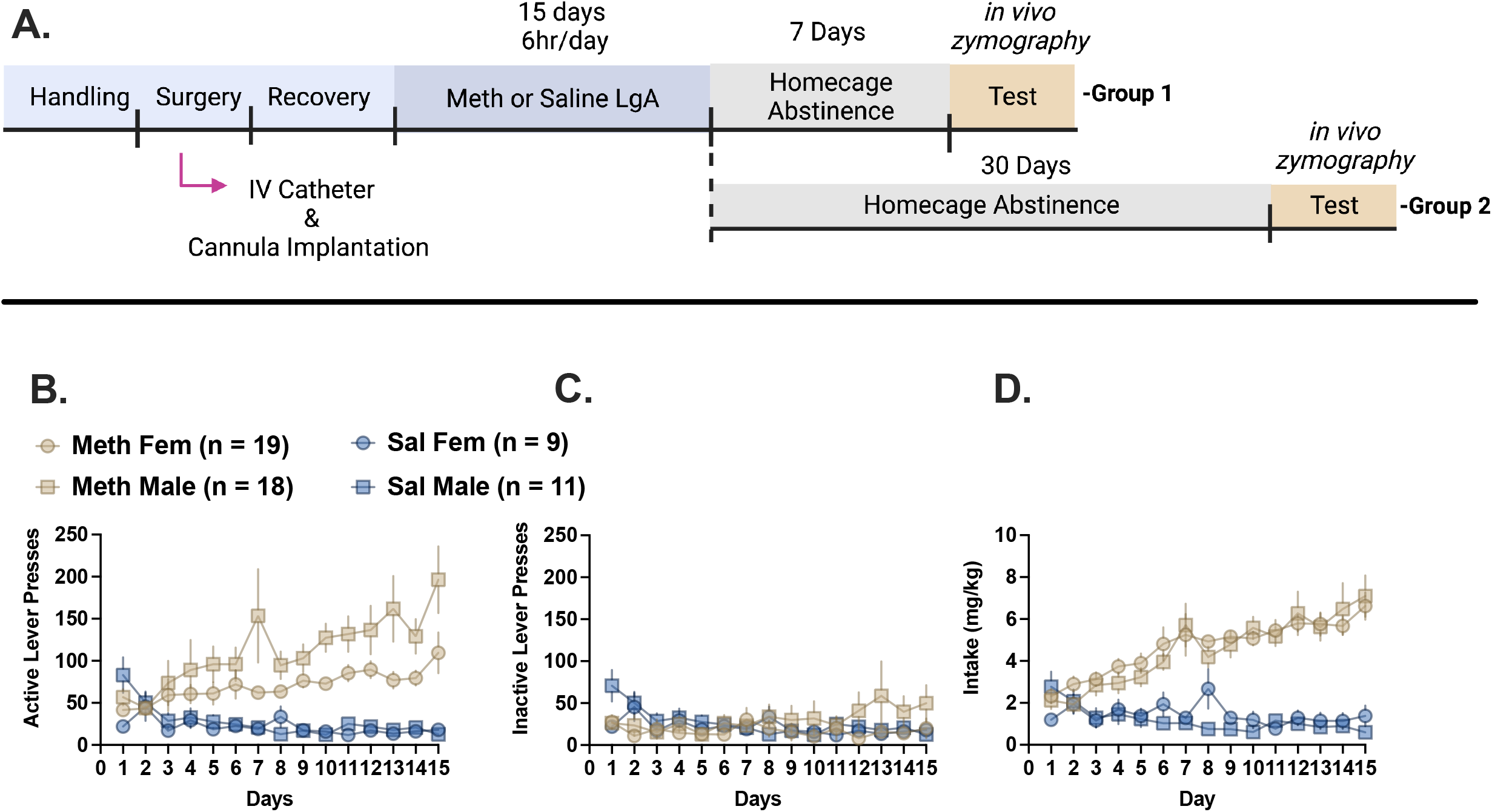
Self-administration for methamphetamine and saline. (A) Experimental timeline for self-administration, home-cage abstinence, *in vivo* zymography and cued-relapse testing. (B) Active lever responding for meth or saline over 15 daily 6-hr sessions in male and female rats. Significance is denoted in the text, in general males responded more on the active lever than females. (C) Inactive lever responding for meth or saline over 15 daily 6-hr in both sexes. There were no significant differences in inactive lever responding. (D) Meth intake between males and females. There are no differences in the amount of meth or saline intake between males and females. Both sexes took more meth than saline.

When adjusted for mg/kg body weight, male and female rats had similar meth and saline intake. The main effect of sex was not significant, nor did sex interact with any other variable. Meth intake increased over time, whereas saline intake did not [day x drug interaction, *F* (14, 734) = 10.06, *p* < 0.0001]. No other interactions were significant, but the main effects of day and drug reached significance [day, *F* (14, 734) = 4.46, *p* < 0.0001; drug, *F* (1, 53) = 85.3, *p* < 0.0001].

### 3.2 MMPs are Involved in Cued Relapse to Methamphetamine During Abstinence

Early and late abstinence were modeled via 7 or 30 days of home cage abstinence followed by cued-relapse testing. On the cued relapse test (Fig 2A), there was an interaction between drug group (saline cue, meth cue day 7, and meth cue day 30) and lever (active and inactive) [*F* (2, 30) = 5.967, *p* > 0.0066]. The main effects of drug [*F* (2, 30) = 16.90, *p* > 0.0001] and lever [*F* (1, 30) = 20.98, *p* > 0.0001] were also significant. After 7 and 30 days of home cage abstinence, meth rats responded more on the active lever relative to saline [Holm-Sidak day 7, *p*<0.0011; day 30 *p*<0.001] and responding was higher after 30 days of abstinence relative to 7 days [*p*<0.01], indicating that the seeking response to meth cues had incubated over increasing withdrawal time. There were no differences in responding on the inactive lever between groups.

**Figure 2.**
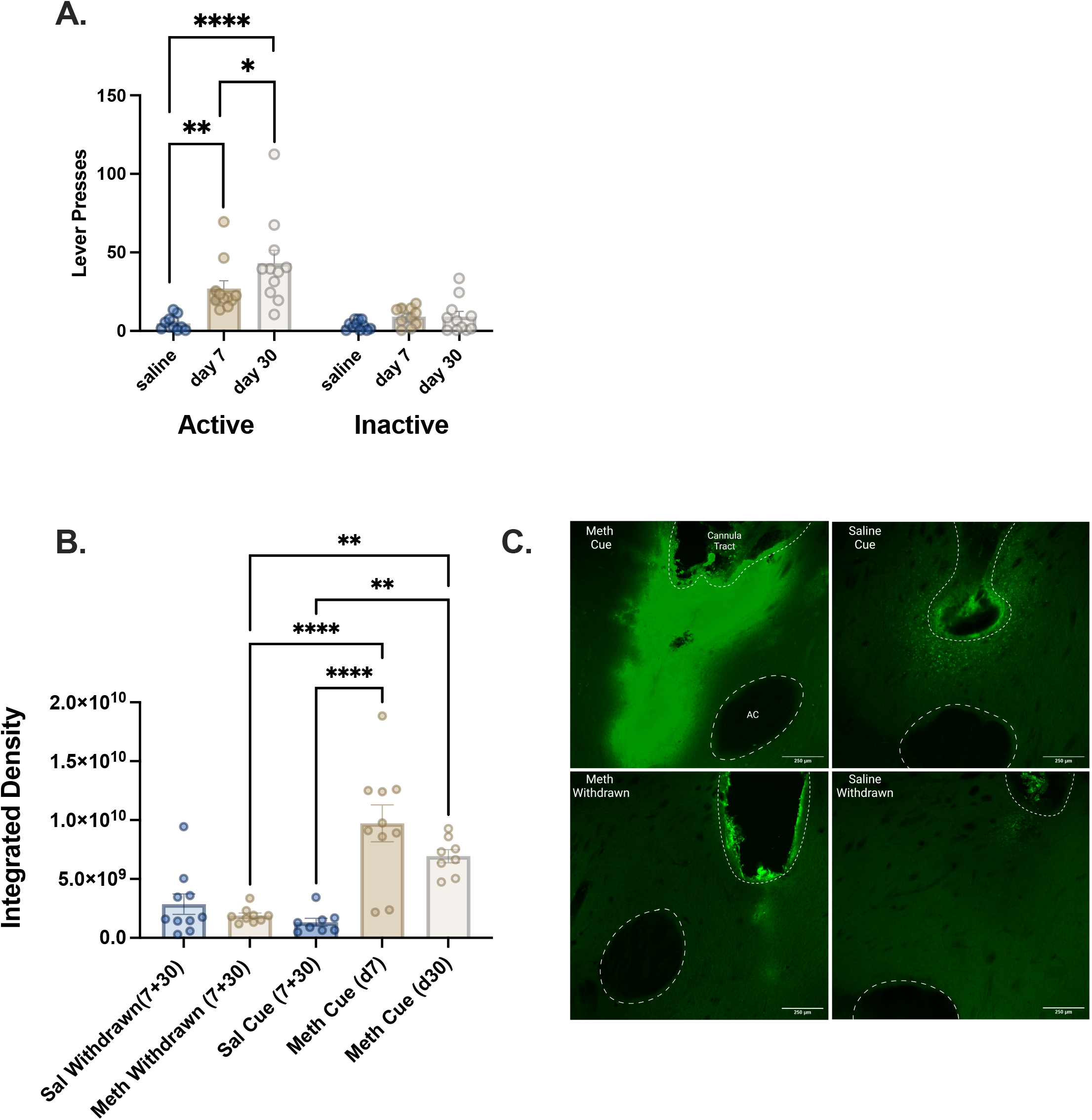
Incubation of meth seeking and cue-induced MMP-2,9 activity. (A) Active and inactive lever pressing during cued-relapse testing. Active lever pressing was significantly increased after 7 and 30 days of home cage abstinence. Further, active lever pressing on abstinence day 10 was elevated relative to day 7, indicating an incubation of meth seeking over time. There were no changes in inactive lever responding. (B) MMP-2,9 activity as measured by integrated density. MMP-2,9 activity was increased between meth-withdrawn (no behavioral test) and cued relapse testing after 7 or 30 days of home cage abstinence. MMP-2,9 activity is also significantly increased between the saline cue group and both meth cue groups. No increase in MMP-2,9 activity was found between the two meth cue groups. (C) Representative micrograph of in vivo zymography assay in the nucleus accumbens core (10x). White dotted lines outline the cannula tract and anterior commissure. Scale bar = 250 μM. * = *p* < 0.05, ** = p < 0.001, and **** = p < 0.0001

**Figure 3.**
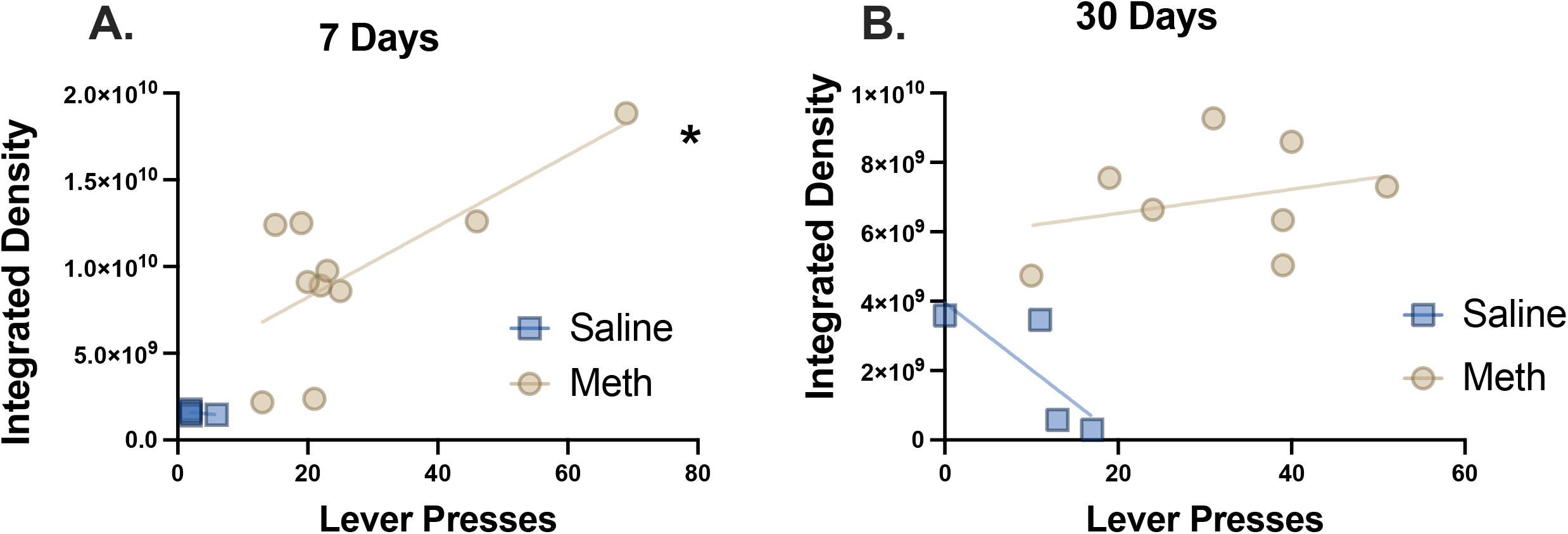
MMP-2,9 activity correlated with active lever pressing during cued-relapse testing. (A) During early abstinence, day 7, MMP-2,9 activity positively correlated with active lever pressing, indicating that the more cued responses presented the higher the integrated density MMP-2,9. (B) During late abstinence, day 30, lever responding and MMP-2,9 integrated density was not correlated. * = *p* < 0.05

Analysis of MMP-2,9 activity via integrated density (Fig 2B) also included meth and saline “withdrawn” groups that underwent self-administration behavior and were not placed back into the behavioral chamber. These groups account for MMP activation without experiencing an associative cue. There was a significant one-way ANOVA [*F* (4, 40) = 15.53, *p* > 0.0001] demonstrating MMP-2,9 activity in the NAcore was significantly increased in the meth cue groups on abstinence day 7 (*p*<0.0001) and day 30 (*p*<0.004) relative to the meth withdrawn group. Additionally, these two meth cue groups had higher MMP-2,9 activity relative to the saline cue group suggesting that exposure to a drug-paired cue is necessary to activate MMP-2,9 in the NAcore (day 7, *p*<0.0001; day 30 *p*<0.0019). MMP-2,9 activity was not different between the two meth-cue timepoints (*p*=0.158). We hypothesized that MMP activity would increase with incubation of drug craving and increased lever pressing during relapse. While both meth abstinent groups differed from saline controls, there was no statistical difference was found between the two meth cue groups (7 or 30 days of home cage abstinence), indicating that MMP-2,9 activity did not parallel incubated behavior. MMP-2,9 activity in the NAcore between both withdrawn groups (saline and meth) and the saline cue group were not statistically different.

### 3.4 MMP-2,9 Activity and Active Lever Pressing During Relapse are Correlated at 7, but not 30 Days of Home Cage Abstinence

Previous studies examining cue-induced cocaine reinstatement in rats extinguished from cocaine self-administration found that the intensity of MMP-2,9 induced fluorescence was correlated with reinstated active lever pressing. To examine whether MMP activity was predictive of behavior in our model of meth use and relapse, we ran correlational analyses of between lever pressing during the cued-relapse test for meth and saline and integrative density of MMP-2,9 activity. Consistent with previous cocaine studies, we observed a significant relationship between active lever pressing and MMP-2,9 activity overall (*r*=0.296, *p*=0.0239). This relationship was carried by data from the 7-day meth abstinent group (*r*=0.507, *p*=0.021) with the 30-day group showing no correlation (r=0.085, p=0.48). No relationships were found between the saline-cue group and MMP activity at any timepoint.

## 4.0 Discussion

The current experiments demonstrated that MMP-2,9 is elevated during cued meth seeking after 7 and 30 days of abstinence. Lever pressing increased after 30 withdrawal days relative to the earlier time point indicating incubation of meth cues; however, MMP-2,9 activity did not become greater after 30 days of withdrawal compared to 7 days. Moreover, MMP-2,9 activity was correlated with lever pressing only in the 7-day withdrawal group, not the 30-day group. Together, these data are consistent with previous studies showing that presenting cocaine-, nicotine-or heroin-associated cues induces MMP-2,9 activity^24– 26^ and the likelihood that MMP-2,9 activity may be necessary for meth seeking, akin to what was shown previously with cocaine, cannabis and heroin using small molecule inhibitors of MMP-2 and/or MMP-9 activity^25,26,30,31^.

### Sex differences

Females are reported to acquire meth self-administration faster than males^32^, take more drug^11,13,32^, and have greater motivation to seek the drug^29,32,33^. However, sex differences are not always identified, which can be accounted for by session length, drug dose administered, or FR value^27,34,35^. In the current study, we used a 6 hr meth protocol for 15 days of self-administration both sexes and while we found sex differences, it was males, not females that pressed the active lever more. This finding replicates our previous study using the same meth regimen^35^. We posit that animals titrate their meth infusions because overall intake between the two sexes was not different^35^. Additionally, consistent with our previous observation with cue-induced meth seeking after extinction training^27^, we found no sex differences in our cued meth seeking task.

### Meth cue-induced increase in MMP-2,9 activity

The induction of MMP-2,9 activity is necessary with classic forms of neuronal plasticity, such as long-term potentiation in the hippocampus^21^, as well as certain learning and memory functions^36,37^, including memory reconsolidation^30^. Because of meth’s high potential for relapse, we had hypothesized that MMP-2,9 activity would be increased during cued-meth relapse following abstinence. Indeed, MMP-2,9 activity was increased after 7 or 30 days of home cage abstinence. Thus, cue-induced meth seeking is similar to cannabis^31^, cocaine^26^, heroin^38^, and nicotine^26^ where drug cues induce parallel increases in behavioral seeking and MMP-2,9 activity. This indicates that in the NAcore synaptic remodeling initiated by MMP-2,9 activity is a shared feature of cued seeking across many addictive drugs. Moreover, while not examined in the present study, cue-induced sucrose seeking does not increase NAcore MMP-2,9 activity^26,38^, indicating that this is a feature of addictive drugs not biological reinforcers. Also, all the previous studies were conducted after withdrawal plus extinction training, except for one study examining heroin seeking that was conducted using both an abstinent and extinction withdrawal model^38^. These authors observed that 2 weeks of heroin abstinence was not associated with increased MMP-2,9 activity, but that extinguished rats show constitutive increases selectively in MMP-2 only around dendrites of NAcore medium spiny neurons expressing D2-dopamine receptors, not around those expressing D1-receptors [D2- and D1-medium spiny neurons (MSNs), respectively]. Akin to the heroin study, we found that meth abstinence without cued seeking (withdrawn group) was not associated with constitutive increases in MMP-2,9 activity.

### Association between incubated meth seeking and MMP-2,9 activity during early abstinence

We also hypothesized that akin to the behavioral incubation of meth seeking with increasing withdrawal period that MMP-2,9 activity would also increase. This latter hypothesis was not borne out by our data that showed equivalent MMP-2,9 activity in NAcore between the 7-day and 30-day meth cue groups. Moreover, while MMP-2,9 activity and cue-induced active lever pressing were linearly correlated after 7 days of withdrawal, the correlation collapsed by 30 days of withdrawal. The correlation with behavior on day 7 is akin to previous studies with cocaine^26,39^ that showed correlations after 2-3 weeks of extinction training during withdrawal. The fact that behavioral meth seeking increased from day 7 to day 30 of withdrawal but MMP-2,9 activity did not has several potential implications and explanations.

It is possible that incubated seeking is not correlated with MMP-2,9 activity because cue-induced seeking, with is shared on day 7 and 30 of withdrawal, has a distinct molecular substrate from the incubation process which serves to augment drug seeking but does not initiate drug seeking. Thus, incubation may be mediated through a different signaling mechanism within the NAcore, or via different circuitry mechanisms altogether. For example, incubation of abstinent cocaine seeking is mediated by a constitutive increase in calcium permeable (CP) AMPA receptors in the accumbens^40^, which is separate from the synaptic potentiation mediated by cue-induced glutamate spillover and activation of MMP-2,9 to stimulate integrin receptors and promote non CP-AMPA receptor insertion and spine head enlargement^41^. Similar to cocaine, meth self-administration and abstinent withdrawal is associated behavioral incubation increases CP-AMPA receptors in the NAcore^42^. Thus, it may be a combination of MMP-2,9 initiated seeking and MMP-2,9 independent incubation of seeking that resulted in the failed correlation between MMP-2,9 and behavior in incubated meth seeking since individual differences in the extent of incubated responding would obscure the effect of seeking without incubation on day 7.

Alternatively, a previous study examining cell-specific MMP-2,9 activation by heroin cue-induced seeking found that activity increased around D1-, but was reduced around D2-MSNs^38^. It is possible that recruitment of distinct MMP-2,9 mechanisms around different cell types after 30 days of withdrawal could negate the correlation with the cell nonspecific measure used in the present study. Also, these authors found that while MMP-9 activity was elevated by cues around D1-MSNs, it was MMP-2 activity that decreased around D2-MSNs. In future studies it is possible that discerning between different MMPs would reveal a more causal relationship between incubated meth seeking and MMP activity. Finally, there are distinctions between meth and most other addictive drugs in terms of how cues affect tetrapartite plasticity in the NAcore. Notably, meth self-administration does not alter protein levels of GLT-1 or influence glutamate uptake^43^, which is in contrast with cocaine or heroin^44,45^. Additionally, at the circuit level, inhibition of the PL-NAcore circuit reduces the reinstatement of cocaine seeking following extinction training^46,47^, while inhibition of the IL-NAshell circuit reinstates seeking previously extinguished rats^48,49^. Conversely, while inactivation of the PL-NAcore circuit also reduces cued-seeking in meth, it does not does induce reinstatement of drug seeking when the IL-NAshell circuit is activated^27^. Finally, meth is a well-established neurotoxin when used in high dose, long access regimens^14^ and such an insult to brain function could alter circuit involvement making the NAcore a less critical locus in regulating the incubation of meth seeking. In our model, one or more of these differences could explain the lack of a strong relationship between active lever pressing and MMP activity after 30 days of home cage abstinence.

## Conclusions

MMP-2,9 activation occurs during synaptic plasticity. Cue-induced drug seeking, including meth seeking, induces transient plasticity and we show that MMP-2,9 activity is also elevated during cued meth seeking. Our data also indicate that the incubation of meth seeking produced by long periods of abstinence may result from different mechanisms than MMP-2,9 dependent drug seeking. However, we propose that the cue initiating incubated seeking requires MMP-2,9 activity and testing this possibility is the topic of future studies. If proven true, it marks modulation of MMP-2,9 activity as a candidate mechanism for inhibiting meth craving and seeking.

## Acknowledgements

Methamphetamine HCL was provided by the National Institute on Drug Abuse drug supply program (Bethesda, MD, USA). We thank Dr. Constanza Garcia-Keller for technical assistance.

## Author Contributions

SIL, PWK, and CR designed the experiments. SIL executed the experiments. RH provided technical expertise. SIL, JC, SW and KN assisted in data collection. SIL wrote the manuscript. PWK and CR edited the final manuscript.

